# Large-scale pathway-specific polygenic risk, transcriptomic community networks and functional inferences in Parkinson disease

**DOI:** 10.1101/2020.05.05.079228

**Authors:** S Bandres-Ciga, S Saez-Atienzar, JJ Kim, MB Makarious, F Faghri, M Diez-Fairen, H Iwaki, H Leonard, J Botia, M Ryten, D Hernandez, JR Gibbs, J Ding, Z Gan-Or, A Noyce, L Pihlstrom, A Torkamani, SW Scholz, B Traynor, D Ehrlich, CR Scherzer, M Bookman, M Cookson, C Blauwendraat, MA Nalls, AB Singleton, on behalf of the International Parkinson Disease Genomics Consortium

## Abstract

Polygenic inheritance plays a central role in Parkinson disease (PD). A priority in elucidating PD etiology lies in defining the biological basis of genetic risk. Unraveling how risk leads to disruption will yield disease-modifying therapeutic targets that may be effective. Here, we utilized a high-throughput and hypothesis-free approach to determine biological pathways underlying PD using the largest currently available cohorts of genetic data and gene expression data from International Parkinson’s Disease Genetics Consortium (IPDGC) and the Accelerating Medicines Partnership - Parkinson’s disease initiative (AMP-PD), among other sources. We placed these insights into a cellular context. We applied large-scale pathway-specific polygenic risk score (PRS) analyses to assess the role of common variation on PD risk in a cohort of 457,110 individuals by focusing on a compilation of 2,199 publicly annotated gene sets representative of curated pathways, of which we nominate 46 pathways associated with PD risk. We assessed the impact of rare variation on PD risk in an independent cohort of whole-genome sequencing data, including 4,331 individuals. We explored enrichment linked to expression cell specificity patterns using single-cell gene expression data and demonstrated a significant risk pattern for adult dopaminergic neurons, serotonergic neurons, and radial glia. Subsequently, we created a novel way of building *de novo* pathways by constructing a network expression community map using transcriptomic data derived from the blood of 1,612 PD patients, which revealed 54 connecting networks associated with PD. Our analyses highlight several promising pathways and genes for functional prioritization and provide a cellular context in which such work should be done.

## INTRODUCTION

Although a great deal of progress in understanding the genetic underpinnings of familial and sporadic Parkinson disease (PD) has been made, the biological basis and cellular context of this risk remains unclear. We have learned that about 1-2 % of PD is associated with a classical Mendelian inheritance pattern, while the majority of disease is driven by a complex set of factors in which polygenic risk seems to play a crucial role [4]. The fact that many of the genes that contain disease-causing mutations also map within risk loci identified by genome-wide association studies (GWAS), supports the notion that common pathways are involved in both forms, and therefore these pleomorphic genes might interact to regulate downstream common targets in both monogenic and non-monogenic PD [22].

Several common molecular processes have been suggested as critical in PD pathophysiology, including lysosome mediated autophagy, mitochondrial dysfunction, endosomal protein sorting and recycling, immune response, and synaptic transmission [3]. A goal in much of this work has been to unify the proteins encoded by PD-linked genes into common pathways. For instance, some success has been seen in this regard within the autosomal recessive genes *PINK1, PRKN*, and *DJ-1*, which share a common cellular mechanism: mitochondrial quality control and regulation [10][16]. However, despite this success, the PD genetics field is still facing the challenge of understanding how genetic risk variants may disrupt biological processes and drive the underlying pathobiology of the disease. In the current era, using genetics to understand the disease process is a key milestone to facilitate the development of targeted therapies.

A priority in elucidating PD etiology lies in defining cumulative risk. GWAS continues to expand the number of genes and loci associated with disease [15], but the majority of these contributors individually exert small effects on PD risk. Current estimates of heritability explained by GWAS loci suggest that there is still an important component of risk yet to be discovered.

Here we present a novel high-throughput and hypothesis-free approach to detect the existence of PD genetic risk linked to any particular biological pathway. We apply polygenic risk score (PRS) to a total of 2,199 curated and well-defined gene sets representative of canonical pathways publicly available in the Molecular Signature Database v7.2 (MSigDB)[24] to define the cumulative effect of pathway-specific genetic variation on PD risk. To assess the impact of rare variation on PD risk explained by significant pathways, we perform gene-set burden analyses in an independent cohort of whole-genome sequencing (WGS) data, including 2,101 cases and 2,230 controls.

Additionally, we explore cell type expression specificity enrichment linked to PD etiology by using single-cell RNA sequencing data from brain cells. Furthermore, we use graph-based analyses to generate *de novo* pathways that could be involved in disease etiology by constructing a transcriptome map of network communities based on RNA sequencing data derived from blood of 1,612 PD patients and 1,042 healthy subjects.

Subsequently, we perform summary-data based Mendelian randomization (SMR) analyses to prioritize genes from significant pathways by exploring possible genomic associations with expression quantitative trait loci (eQTL) in public databases and nominate overlapping genes within our transcriptome communities for follow-up functional studies. Finally, we present a user-friendly platform for the PD research community that enables easy and interactive access to these results (https://pdgenetics.shinyapps.io/pathwaysbrowser/).

## METHODS

### Gene set selection representative of canonical pathways

The Molecular Signatures Database (MSigDB database v7.2) is a compilation of annotated gene sets from various sources such as online pathway databases, the biomedical literature, and manual curation by domain experts [13, 24]. We selected the collection “Canonical Pathways” composed of 2,199 curated gene sets of pathways annotated from the following databases; Reactome (1,499), KEGG (186), BIOCARTA (289), Pathway Interaction Database (196), Matrisome project (10), Signaling Gateway (8), Sigma Aldrich (10), SuperArray SABiosciences (1) (http://software.broadinstitute.org/gsea/msigdb).

### Genotyping data: Cohort characteristics, quality control procedures and study design

To assess PD risk, summary statistics from Chang et al., 2017 PD GWAS meta-analysis involving 26,035 PD cases and 403,190 controls of European ancestry were used as the *reference dataset* for the primary analysis to define risk allele weights. In this study, there were 7,909,453 imputed SNPs tested for association with PD with a minor allele frequency (MAF) > 0.03. Recruitment and genotyping quality control procedures were described in the original report (Chang et al., 2017). Individual-level genotyping data not included in Chang et al. 2017 and from the last GWAS meta-analysis [15] was then randomly divided as the *training* and *testing datasets*. The *training dataset* used to construct the PRS consisted of 7,218 PD cases and 9,424 controls, while the *testing dataset* to validate the results consisted of 5,429 PD cases and 5,814 controls, all of European ancestry **(see Figure 1 for analysis workflow and rationale summary)**. Demographic and clinical characteristics of the cohorts under study are given in **Supplementary Table 1**.

**Figure 1.**
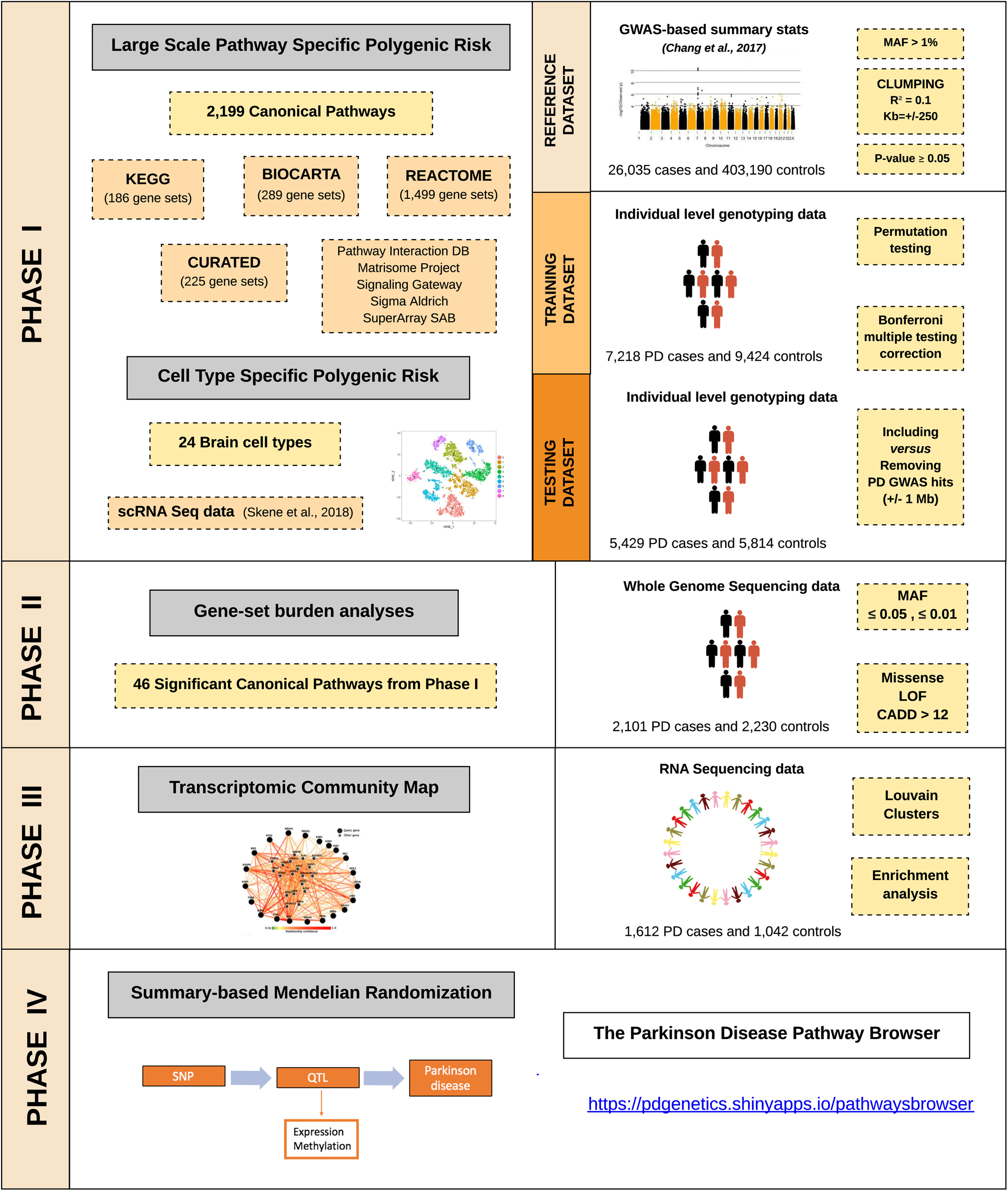
Workflow and rationale summary.

Additional details of these cohorts, along with detailed quality control (QC) methods, can be found in Nalls et al., 2019. For sample QC, in short, individuals with low call rates (< 95%), discordance between genetic and reported sex, heterozygosity outliers (F-statistic cutoff of >−0.15 and <0.15) and ancestry outliers (+/− 6 standard deviations from means of eigenvectors 1 and 2 of the 1000 Genomes phase 3 CEU and TSI populations from principal components) were excluded. Further, for genotype QC, variants with a missingness rate of > 5%, minor allele frequency <0.05, exhibiting Hardy–Weinberg Equilibrium (HWE) <1E-5 and palindromic SNPs were excluded. Remaining samples were imputed using the Haplotype Reference Consortium (HRC) on the University of Michigan imputation server under default settings with Eagle v2.3 phasing based on *Haplotype Reference Consortium r1.1 2016* (http://www.haplotype-reference-consortium.org), and variants with an imputation quality (R^2^ >0.3) were included.

### Polygenic effect scores for individual biological pathways versus PD risk

A polygenic effect score (PES) was generated to estimate polygenic risk for each of the 2,199 gene sets representative of biological pathways and then tested for association with PD. PES was calculated based on the weighted allele dose as implemented in *PRSice2* (v2.1.1) (https://github.com/choishingwan/PRSice)[7]. Using the *reference dataset*, we selected variants with a summary statistic p-value of association less than or equal to 0.05 and with MAF > 1%. We extracted these variants from the *training dataset*, and linkage disequilibrium (LD) clumping was performed using the default r^2^ = 0.1 and 250 Kb of distance. Then, 1,000 permutations of sample labels were implemented to generate association p-value estimates for each pathway. A p-value threshold = 0.05 was considered to prefilter the inclusion of variants in an effort to avoid overfitting when comparing across gene sets as well as improve computational efficiency. The permutation test in the *training dataset* provided a Nagelkerke’s pseudo r^2^ value after adjusting for an estimated prevalence of 0.005 (aged population estimate as per Gasser and colleagues), age at onset for cases and age at examination for controls, gender and 20 PCs to account for population stratification. For those pathways surpassing Bonferroni multiple testing correction (p-value corrected = 0.05/2,199 pathways = 2.27E-5), PES was then tested in an independent cohort (*testing dataset)* in a similar way, and overlapping gene-sets significantly associated with PD risk were reported. In an attempt to explore what biological processes were associated with PD risk after excluding known risk factors, the same analyses were performed after removing the 90 known PD GWAS hits [15] and additional SNPs located 1 Mb upstream and downstream from the signal. PES analyses considered that all the variants conferred risk under the additive model and did not cover regulatory regions adjacent to the up or downstream of the genes or intergenic variants.

### Whole genome sequencing data: Cohort characteristics and quality control procedures

The following eight cohorts were utilized in this study; Biofind (https://biofind.loni.usc.edu/), NABEC [9], LNG Path confirmed, PDBP (https://pdbp.ninds.nih.gov/), NIH PD CLINIC, PPMI (https://www.ppmi-info.org/), WELLDERLY and UKBEC. Clinical and demographic characteristics of the cohorts under study are summarised in **Supplementary Table 2**. Participants included sporadic PD cases clinically diagnosed by experienced neurologists. PD cases met criteria defined by the UK PD Society Brain Bank. This included 2,101 cases and 2,230 controls. All individuals were of European descent and were not age- or gender-matched.

DNA sequencing was performed using two vendors; Macrogen and USUHS. For samples sequenced at Macrogen one microgram of each DNA sample was fragmented by Covaris System and further prepared according to the Illumina TruSeq DNA Sample preparation guide to obtain a final library of 300-400 bp average insert size. Libraries were multiplexed and sequenced on Illumina HiSeq X platform. For samples sequenced by USUHS, DNA samples were processed using the Illumina TruSeq DNA PCRFree Sample Preparation kit, starting with 500 ng input and resulting in an average insert size of 310 bp. USUHS processed single-libraries on single lanes on HiSeq X flowcells, and the Macrogen protocol used multiplexing. Paired-end read sequences were processed in accordance with the pipeline standard developed by the Centers for Common Disease Genomics [35]. The GRCh38DH reference genome was used for alignment as specified in the FE standardized pipeline [36]. The Broad Institute’s implementation of this FE standardized pipeline, which incorporates the GATK (2016) Best Practices is publicly available and used for WGS processing. Single-nucleotide (SNV) and InDel variants were called from the processed WGS data following the GATK (2016) Best Practices [1] using the Broad Institute’s workflow for joint discovery and Variant Quality Score Recalibration (VQSR). For quality control, each sample was checked using common methods for genotypes as well and sequence-related metrics. Using Plink v1.9 [6], each sample’s genotype missingness rate (< 95%), heterozygosity rate (exceeding +/- 0.15 F-stat), and gender were checked. The King v2.1.3 kinship tool (8) was used to check for the presence of duplicate samples. Sequence and alignment related metrics generated by the Broad’s implementation of the FE standardized pipeline were inspected for potential quality problems. This included the sample’s mean sequence depth (< 30X) and contamination rate (> 2%), as reported by VerifyBamID (9), and single nucleotide variant count as reported by Picard’s CollectVariantCallingMetrics (< 3 StDev) based on the sample’s genomic vcf (gvcf). Principal components (PCs) were created for each dataset using PLINK. For the PC calculation, variants were filtered for minor allele frequency (>0.01), genotype missingness (<0.05), and HWE (P =>1E-6), and minor allele count < 3. GCTA [30] was used to remove cryptically related at the level of first cousins or closer (sharing proportionally more than 12.5 % of alleles).

### Gene-set burden analyses

The sequence kernel association test – optimal (SKAT-O) [12] was implemented using default parameters in RVTESTS [32] to determine the difference in the aggregate burden of rare coding genetic variants (minor allele count ≥ 3) between PD cases and controls for the nominated pathways by PRS. SKAT-O was applied to aggregate genetic information across defined genomic regions to test for associations with pathways of interest under two frequency levels (MAF ≤ 0.05 and MAF ≤ 0.01) and two functionality categories (missense, loss of function and Combined Annotation Dependent Depletion (CADD) score > 12 representing between 1-10% predicted most pathogenic variants in the genome). Covariates including gender, age at onset (cases), age at enrollment (controls), and 10 PCs were included to adjust the analyses. ANNOVAR was used for variant annotation [28].

### Network expression community map in gene expression data

Baseline peri-diagnostic RNA sequencing data derived from blood for 1,612 PD patients and 1,042 healthy subjects available from the Parkinson Progression Marker Initiative (PPMI) was used to construct a network of expression communities based on a graph model with Louvain clusters. This cleaned and normalized data was downloaded from the Accelerating Medicines Partnership for Parkinson’s disease (AMP-PD) on March 1^st^, 2020. Library preparation, protocol, and transcriptomic quality control procedures can be found in detail in the original source https://amp-pd.org/transcriptomics-data. Prior to analyses, all data for the baseline visit was extracted. Data for each gene was then z-transformed to a mean of zero and a standard deviation of one. Scikit-learn’s extraTreeClassifier option was used to extract coding gene features for inclusion in the network builds that are likely to contribute to classifying cases versus controls under default settings in the feature selection phase, leaving 8,3k protein-coding genes for candidate networks [20]. Following this feature extraction phase, controls were excluded, and case-only correlations were calculated for all remaining gene features. Next, this correlation structure was converted to a graph object using NetworkX [26]. We filtered for network links at positive correlations (upregulated in cases together) between genes greater than or equal to 0.8. Subsequently, the Louvain algorithm was employed to build network communities within this graph object derived from the selected feature set [2].

Finally, pathway enrichment analysis within expression communities was performed to further dissect its biological function using the function g:GOSt from g:ProfileR [17]. The significance of each pathway was tested by hypergeometric tests with Bonferroni correction to calculate the error rate of each network.

### Cell-type polygenic risk enrichment analysis

Single-cell RNA sequencing data [23] based on a total of 9,970 cells obtained from several mouse brain regions (neocortex, hippocampus, hypothalamus, striatum, and midbrain) was used to explore cell types associated with PD risk. This dataset includes the specificity of expression for each gene within each cell type where values range from zero to one and represent the proportion of the total expression of a gene found in one cell type compared to all cell types. The closer the score is to 1, the more specific is the expression in that particular cell type. PRS R^2^ (variance) was calculated within each cell type quintile, ranging from 0 to 1. If a particular cell type is associated with PD risk, it is expected to observe a shift in the linear distribution with low PRS R^2^ for PD in non-specific gene sets (i.e., lower quintiles) and a higher PRS R^2^ for PD in more specific gene sets (i.e., higher quintiles). Linear regression adjusted by the number of SNPs included in the PRS was performed to assess the trend of increased PRS R^2^ per quintile of cell type expression specificity.

### Summary-data-based Mendelian Randomization Quantitative Trait Loci analyses

Two-sample SMR was applied to explore the enrichment of *cis* eQTLs within the 46 pathways nominated by our large-scale PRS analysis. The methodology can be interpreted as an analysis to test if the effect size of genetic variants influencing PD risk is mediated by gene expression or methylation to prioritize genes underlying these pathways for follow-up functional studies [34]. QTL association summary statistics from well-curated expression datasets were compared to Nalls et al., 2019 summary statistics after extracting the pathway-specific independent SNPs considered as the instrumental variables. Expression datasets used for these analyses include estimates for cis-expression from the Genotype-Tissue Expression (GTEx) Consortium (v6; whole blood and 10 brain regions), the Common Mind Consortium (CMC; dorsolateral prefrontal cortex), the Religious Orders Study and Memory and Aging Project (ROSMAP), and the Brain eQTL Almanac project (Braineac; 10 brain regions). Additionally, we studied expression patterns in blood from the largest eQTL meta-analysis so far [27]. LD pruning and clumping were carried out using default SMR protocols (http://cnsgenomics.com/software/smr). Multi-SMR p-values (gene-level expression summaries for eQTLs) were adjusted by Bonferroni multiple test correction considering the number of genes tested per pathway, and HEIDI was used to detect pleiotropic associations between the expression levels and PD risk that could be biasing the model at a p-value < 0.01 [29]. Effect estimates represent the change in PD odds ratio per one standard deviation increase in gene expression. Enrichment of *cis* expression was assessed per pathway and per tissue. The number of genes tested per pathway were Bonferroni corrected, and a Chisq test was applied to assess whether the proportion of QTLs per pathway was significantly higher than expected by chance.

## RESULTS

### Large scale PES analysis nominates biological pathways involved on PD risk

Out of the 2,199 gene sets representative of biological processes included in this report, 279 gene-sets were significantly associated with PD risk in the *training phase* (Bonferroni threshold for significance 0.05/2,199 pathways = 2.27E-5) (**Supplementary Table 3**, https://pdgenetics.shinyapps.io/pathwaysbrowser/). Following the same analysis workflow, a total of 46 gene sets were replicated in the *testing phase* and nominated as potentially linked to PD risk through common genetic variation **(Table 1, Figure 2A and 2B)**. After excluding the 90 PD risk loci and SNPs located 1 Mb upstream and downstream from the GWAS hits, six gene sets including adaptive immune system, innate immune system, vesicle mediated transport, signaling by G protein-coupled receptors (GPCR) ligand binding, metabolism of lipids and neutrophil degranulation remained significant, suggesting as yet unidentified risk within these pathways (Bonferroni threshold for significance 0.05/2,199 pathways = 2.27E-5) (**Table 2, Figure 2C and 2D**). For an easy interpretation of these findings, significant pathways were clustered in hierarchies according to genetic redundancy, as highlighted in **Supplementary Figure 1-2**.

**Table 1.** Canonical pathways significantly associated with PD risk in the *discovery* and *replication phases* through common variation.

| Gene set | DISCOVERY |  |  |  |  | REPLICATION |  |  |  |  |
| --- | --- | --- | --- | --- | --- | --- | --- | --- | --- | --- |
|  | PRS R2 | Beta | SE | P | Num SNP | PRS R2 | Beta | SE | P | Num SNP |
| ACTIVATION OF AMPK DOWNSTREAM OF NMDARS (REACTOME) | 0.0010 | 0.0936 | 0.0162 | 8.44E-09 | 18 | 0.0014 | 0.1088 | 0.0191 | 1.32E-08 | 9 |
| ADAPTIVE IMMUNE SYSTEM (REACTOME) | 0.0040 | 0.1867 | 0.0164 | 5.67E-30 | 455 | 0.0008 | 0.0815 | 0.0192 | 2.11E-05 | 169 |
| ALPHA SYNUCLEIN PATHWAY (PID) | 0.0015 | 0.1157 | 0.0164 | 1.52E-12 | 34 | 0.0009 | 0.0902 | 0.0192 | 2.54E-06 | 17 |
| ALZHEIMERS DISEASE (KEGG) | 0.0023 | 0.1410 | 0.0163 | 5.91E-18 | 175 | 0.0017 | 0.1208 | 0.0192 | 3.34E-10 | 50 |
| AMYLOID FIBER FORMATION (REACTOME) | 0.0031 | 0.1646 | 0.0164 | 7.60E-24 | 28 | 0.0019 | 0.1286 | 0.0193 | 2.40E-11 | 15 |
| APOPTOTIC CLEAVAGE OF CELLULAR PROTEINS (REACTOME) | 0.0019 | 0.1289 | 0.0164 | 3.71E-15 | 41 | 0.0009 | 0.0890 | 0.0191 | 3.12E-06 | 23 |
| APOPTOTIC EXECUTION PHASE (REACTOME) | 0.0018 | 0.1243 | 0.0164 | 2.91E-14 | 48 | 0.0009 | 0.0867 | 0.0191 | 5.53E-06 | 26 |
| ASPARAGINE N LINKED GLYCOSYLATION (REACTOME) | 0.0020 | 0.1331 | 0.0163 | 3.21E-16 | 235 | 0.0008 | 0.0843 | 0.0191 | 1.01E-05 | 100 |
| CASPASE MEDIATED CLEAVAGE OF CYTOSKELETAL PROTEINS (REACTOME) | 0.0021 | 0.1372 | 0.0164 | 5.91E-17 | 15 | 0.0009 | 0.0900 | 0.0191 | 2.47E-06 | 13 |
| CHROMATIN ORGANIZATION (REACTOME) | 0.0007 | 0.0773 | 0.0163 | 2.11E-06 | 201 | 0.0012 | 0.1004 | 0.0191 | 1.56E-07 | 79 |
| CLASS B 2 SECRETIN FAMILY RECEPTORS (REACTOME) | 0.0015 | 0.1146 | 0.0163 | 1.98E-12 | 65 | 0.0011 | 0.0962 | 0.0191 | 4.84E-07 | 23 |
| CLATHRIN MEDIATED ENDOCYTOSIS (REACTOME) | 0.0013 | 0.1079 | 0.0162 | 3.06E-11 | 151 | 0.0008 | 0.0810 | 0.0191 | 2.22E-05 | 81 |
| COPI DEPENDENT GOLGI TO ER RETROGRADE TRAFFIC (REACTOME) | 0.0016 | 0.1187 | 0.0162 | 2.52E-13 | 76 | 0.0012 | 0.1008 | 0.0191 | 1.38E-07 | 30 |
| COPI MEDIATED ANTEROGRADE TRANSPORT (REACTOME) | 0.0008 | 0.0832 | 0.0162 | 3.00E-07 | 81 | 0.0015 | 0.1132 | 0.0191 | 3.23E-09 | 36 |
| COPII MEDIATED VESICLE TRANSPORT (REACTOME) | 0.0013 | 0.1070 | 0.0163 | 5.69E-11 | 46 | 0.0020 | 0.1325 | 0.0192 | 5.37E-12 | 20 |
| ER TO GOLGI ANTEROGRADE TRANSPORT (REACTOME) | 0.0012 | 0.1008 | 0.0163 | 5.87E-10 | 114 | 0.0017 | 0.1203 | 0.0191 | 3.25E-10 | 48 |
| GLUTAMATE BINDING ACTIVATION OF AMPA RECEPTORS AND SYNAPTIC PLASTICITY (REACTOME) | 0.0020 | 0.1308 | 0.0163 | 9.99E-16 | 48 | 0.0018 | 0.1252 | 0.0192 | 7.34E-11 | 12 |
| GOLGI ASSOCIATED VESICLE BIOGENESIS (REACTOME) | 0.0014 | 0.1118 | 0.0162 | 4.87E-12 | 61 | 0.0016 | 0.1151 | 0.0191 | 1.79E-09 | 32 |
| GPCR LIGAND BINDING (REACTOME) | 0.0017 | 0.1215 | 0.0163 | 9.49E-14 | 263 | 0.0010 | 0.0915 | 0.0191 | 1.65E-06 | 88 |
| INNATE IMMUNE SYSTEM (REACTOME) | 0.0037 | 0.1790 | 0.0164 | 7.05E-28 | 677 | 0.0009 | 0.0870 | 0.0192 | 6.11E-06 | 281 |
| INTRA GOLGI AND RETROGRADE GOLGI TO ER TRAFFIC (REACTOME) | 0.0019 | 0.1275 | 0.0163 | 4.52E-15 | 156 | 0.0011 | 0.0992 | 0.0191 | 2.19E-07 | 60 |
| INTRA GOLGI TRAFFIC (REACTOME) | 0.0013 | 0.1049 | 0.0163 | 1.13E-10 | 36 | 0.0027 | 0.1526 | 0.0192 | 2.05E-15 | 12 |
| LKB1 PATHWAY (PID) | 0.0011 | 0.0988 | 0.0162 | 1.12E-09 | 40 | 0.0012 | 0.1035 | 0.0191 | 6.25E-08 | 21 |
| LONG TERM DEPRESSION (KEGG) | 0.0017 | 0.1204 | 0.0163 | 1.35E-13 | 170 | 0.0010 | 0.0923 | 0.0192 | 1.47E-06 | 41 |
| LYSOSOME (KEGG) | 0.0009 | 0.0890 | 0.0164 | 5.51E-08 | 85 | 0.0010 | 0.0919 | 0.0192 | 1.67E-06 | 46 |
| MAPK SIGNALING PATHWAY (KEGG) | 0.0013 | 0.1058 | 0.0163 | 8.31E-11 | 322 | 0.0008 | 0.0851 | 0.0191 | 8.34E-06 | 84 |
| METABOLISM OF LIPIDS (REACTOME) | 0.0029 | 0.1585 | 0.0163 | 2.99E-22 | 607 | 0.0009 | 0.0888 | 0.0192 | 3.58E-06 | 227 |
| METABOLISM OF VITAMINS AND COFACTORS (REACTOME) | 0.0011 | 0.0965 | 0.0162 | 2.67E-09 | 201 | 0.0009 | 0.0891 | 0.0191 | 3.00E-06 | 86 |
| METABOLISM OF WATER SOLUBLE VITAMINS AND COFACTORS (REACTOME) | 0.0009 | 0.0881 | 0.0162 | 5.47E-08 | 112 | 0.0008 | 0.0829 | 0.0191 | 1.38E-05 | 53 |
| NEUROACTIVE LIGAND RECEPTOR INTERACTION (KEGG) | 0.0020 | 0.1316 | 0.0163 | 6.58E-16 | 328 | 0.0011 | 0.0988 | 0.0191 | 2.34E-07 | 91 |
| NEURONAL SYSTEM (REACTOME) | 0.0044 | 0.1961 | 0.0164 | 7.03E-33 | 880 | 0.0012 | 0.0994 | 0.0191 | 1.92E-07 | 191 |
| NEUROTRANSMITTER RECEPTORS AND POSTSYNAPTIC SIGNAL TRANSMISSION (REACTOME) | 0.0023 | 0.1430 | 0.0163 | 1.76E-18 | 390 | 0.0014 | 0.1079 | 0.0191 | 1.65E-08 | 98 |
| NEUTROPHIL DEGRANULATION (REACTOME) | 0.0020 | 0.1316 | 0.0163 | 6.89E-16 | 294 | 0.0008 | 0.0847 | 0.0192 | 1.01E-05 | 128 |
| P38 GAMMA DELTA PATHWAY (PID) | 0.0020 | 0.1334 | 0.0164 | 3.34E-16 | 13 | 0.0012 | 0.1035 | 0.0191 | 6.26E-08 | 11 |
| PARKINSONS DISEASE (KEGG) | 0.0029 | 0.1635 | 0.0168 | 2.09E-22 | 39 | 0.0023 | 0.1398 | 0.0192 | 3.01E-13 | 18 |
| POST TRANSLATIONAL PROTEIN MODIFICATION (REACTOME) | 0.0074 | 0.2568 | 0.0166 | 3.27E-54 | 1165 | 0.0011 | 0.0990 | 0.0191 | 2.20E-07 | 391 |
| PTK6 PROMOTES HIF1A STABILIZATION (REACTOME) | 0.0025 | 0.1580 | 0.0178 | 6.51E-19 | 18 | 0.0009 | 0.0897 | 0.0191 | 2.60E-06 | 13 |
| RETROGRADE TRANSPORT AT THE TRANS GOLGI NETWORK (REACTOME) | 0.0008 | 0.0856 | 0.0163 | 1.52E-07 | 41 | 0.0008 | 0.0817 | 0.0192 | 1.98E-05 | 17 |
| SIGNALING BY GPCR (REACTOME) | 0.0027 | 0.1534 | 0.0164 | 7.91E-21 | 884 | 0.0010 | 0.0947 | 0.0191 | 7.58E-07 | 282 |
| SNARE INTERACTIONS IN VESICULAR TRANSPORT (KEGG) | 0.0008 | 0.0834 | 0.0162 | 2.64E-07 | 29 | 0.0009 | 0.0860 | 0.0191 | 6.67E-06 | 8 |
| TRAFFICKING OF GLUR2 CONTAINING AMPA RECEPTORS (REACTOME) | 0.0016 | 0.1184 | 0.0163 | 3.61E-13 | 37 | 0.0018 | 0.1249 | 0.0192 | 8.29E-11 | 10 |
| TRANS GOLGI NETWORK VESICLE BUDDING (REACTOME) | 0.0015 | 0.1148 | 0.0162 | 1.31E-12 | 76 | 0.0016 | 0.1152 | 0.0191 | 1.79E-09 | 37 |
| TRANSMISSION ACROSS CHEMICAL SYNAPSES (REACTOME) | 0.0030 | 0.1634 | 0.0163 | 1.56E-23 | 517 | 0.0013 | 0.1055 | 0.0191 | 3.31E-08 | 124 |
| TRANSPORT TO THE GOLGI AND SUBSEQUENT MODIFICATION (REACTOME) | 0.0018 | 0.1238 | 0.0163 | 3.14E-14 | 154 | 0.0016 | 0.1184 | 0.0191 | 6.02E-10 | 64 |
| VASOPRESSIN REGULATED WATER REABSORPTION (KEGG) | 0.0012 | 0.1045 | 0.0163 | 1.43E-10 | 55 | 0.0016 | 0.1171 | 0.0192 | 1.03E-09 | 16 |
| VESICLE MEDIATED TRANSPORT (REACTOME) | 0.0044 | 0.1964 | 0.0164 | 5.90E-33 | 577 | 0.0018 | 0.1226 | 0.0192 | 1.57E-10 | 227 |

PRS R2, variance explained by polygenic risk score; SE, standard error; P, P-value; num\_SNP, Number of SNPs included in each pathway analysis

**Figure 2.**
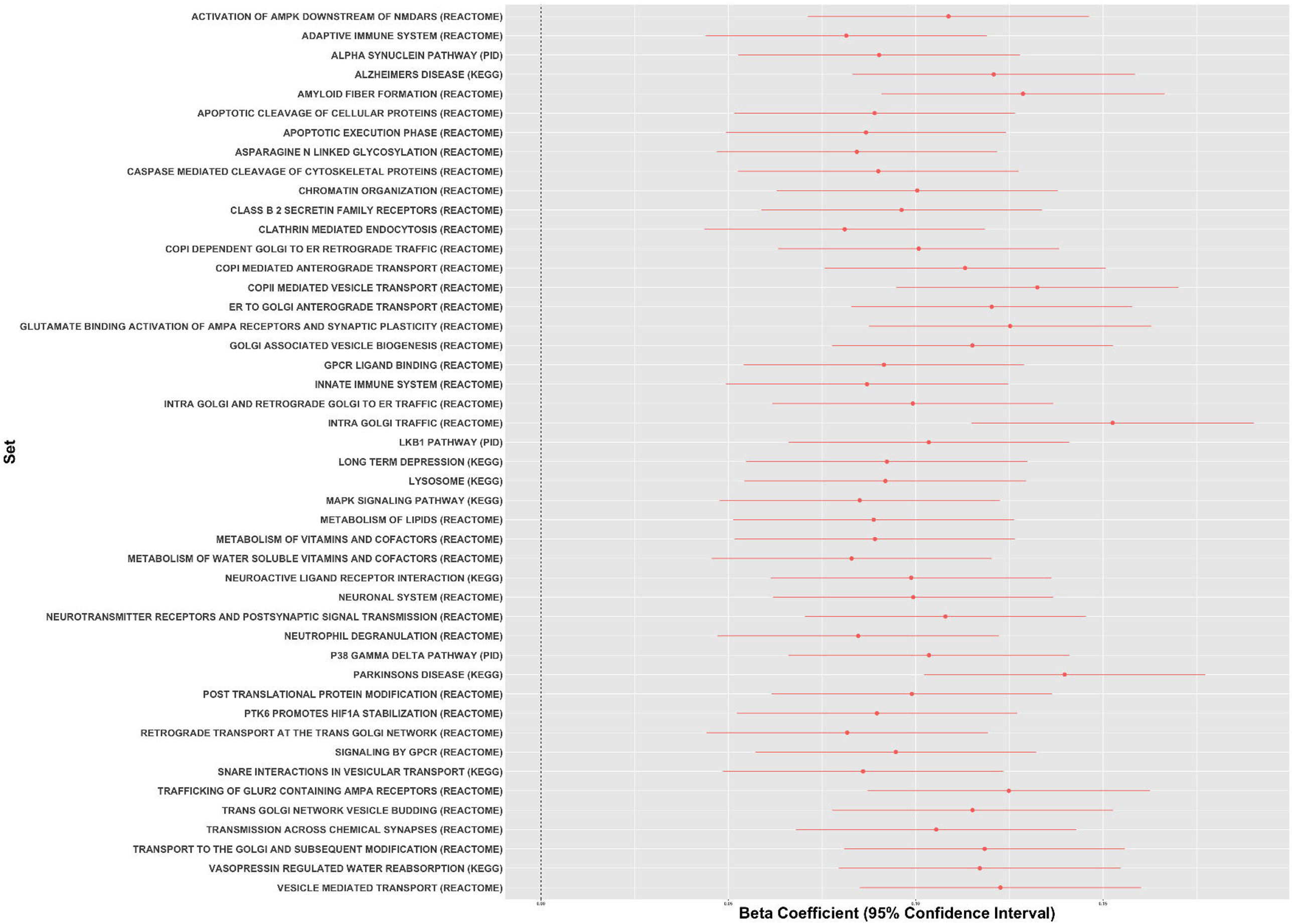
Canonical pathways associated with PD risk through common genetic variation based on PES analyses. Forest plots showing polygenic risk score estimates for the significant canonical pathways in the replication phase including (A) and removing (B) PD known risk loci +/- 1Mb upstream and downstream. Estimates of variance explained by PRS for the significant canonical pathways including () or excluding (D) PD known risk loci +/- 1Mb upstream and downstream.

### Gene-set based burden analyses identify pathways involved on PD risk through rare variation

To test whether the same biological pathways are enriched by rare coding variants, we implemented gene-set based SKAT-O in a large WGS cohort composed of 2,101 PD cases and 2,230 controls. Out of the 46 pathways significantly associated with PD risk through common variation, 18 were linked through low frequency genetic variation (MAF ≤ 5%) and 19 through rare variation (MAF ≤ 1%), at a p-value < 0.05 **(Table 3).** At a MAF threshold ≤ 5%, 12 pathways remained significantly associated with PD risk when focusing only on missense mutations, four when considering only loss of function variants and five when filtering by CADD score > 12 (∼ among the 1-10% most pathogenic variants in the genome) **(Table 3)**. At a MAF threshold ≤ 1%, 12 pathways remained significantly associated with PD risk when focusing only on missense mutations, four when considering only loss of function variants and five when filtering by CADD score > 12 **(Table 3)**. Considering a more stringent p-value (Bonferroni threshold for significance 0.05/46 pathways = 0.001), four gene sets including Alzheimer’s disease, Parkinson’s disease, Neuroactive ligand receptor interaction and GPCR ligand binding remained significant at MAF ≤ 5%. When focusing on MAF ≤ 1%, the four above mentioned gene sets in addition to Aspargine-N-glycosylation were significantly associated with PD risk.

**Table 2.** Canonical pathways significantly associated with PD risk through common variation in the *discovery* and *replication phases* after excluding PD known risk loci +/- 1Mb upstream and downstream

| Gene set | DISCOVERY |  |  |  |  | REPLICATION |  |  |  |  |
| --- | --- | --- | --- | --- | --- | --- | --- | --- | --- | --- |
|  | PRS R2 | Beta | SE | P | Num SNP | PRS R2 | Beta | SE | P | Num SNP |
| ADAPTIVE IMMUNE SYSTEM (REACTOME) | 0.0028 | 0.1560 | 0.0163 | 1.42E-21 | 397 | 0.0002 | 0.0438 | 0.0190 | 2.15E-02 | 182 |
| INNATE IMMUNE SYSTEM (REACTOME) | 0.0026 | 0.1522 | 0.0163 | 1.12E-20 | 621 | 0.0003 | 0.0475 | 0.0192 | 1.34E-02 | 332 |
| VESICLE MEDIATED TRANSPORT (REACTOME) | 0.0022 | 0.1398 | 0.0163 | 1.03E-17 | 515 | 0.0002 | 0.0439 | 0.0190 | 2.12E-02 | 259 |
| SIGNALING BY GPCR (REACTOME) | 0.0019 | 0.1299 | 0.0163 | 1.86E-15 | 816 | 0.0002 | 0.0460 | 0.0190 | 1.58E-02 | 329 |
| METABOLISM OF LIPIDS (REACTOME) | 0.0018 | 0.1265 | 0.0163 | 8.08E-15 | 538 | 0.0004 | 0.0547 | 0.0190 | 4.08E-03 | 275 |
| NEUTROPHIL DEGRANULATION (REACTOME) | 0.0012 | 0.1026 | 0.0163 | 2.73E-10 | 259 | 0.0004 | 0.0593 | 0.0192 | 1.96E-03 | 143 |

PRS R2, variance explained by polygenic risk score; SE, standard error; P, P-value; num\_SNP, Number of SNPs included in each pathway analysis

**Table 3.**
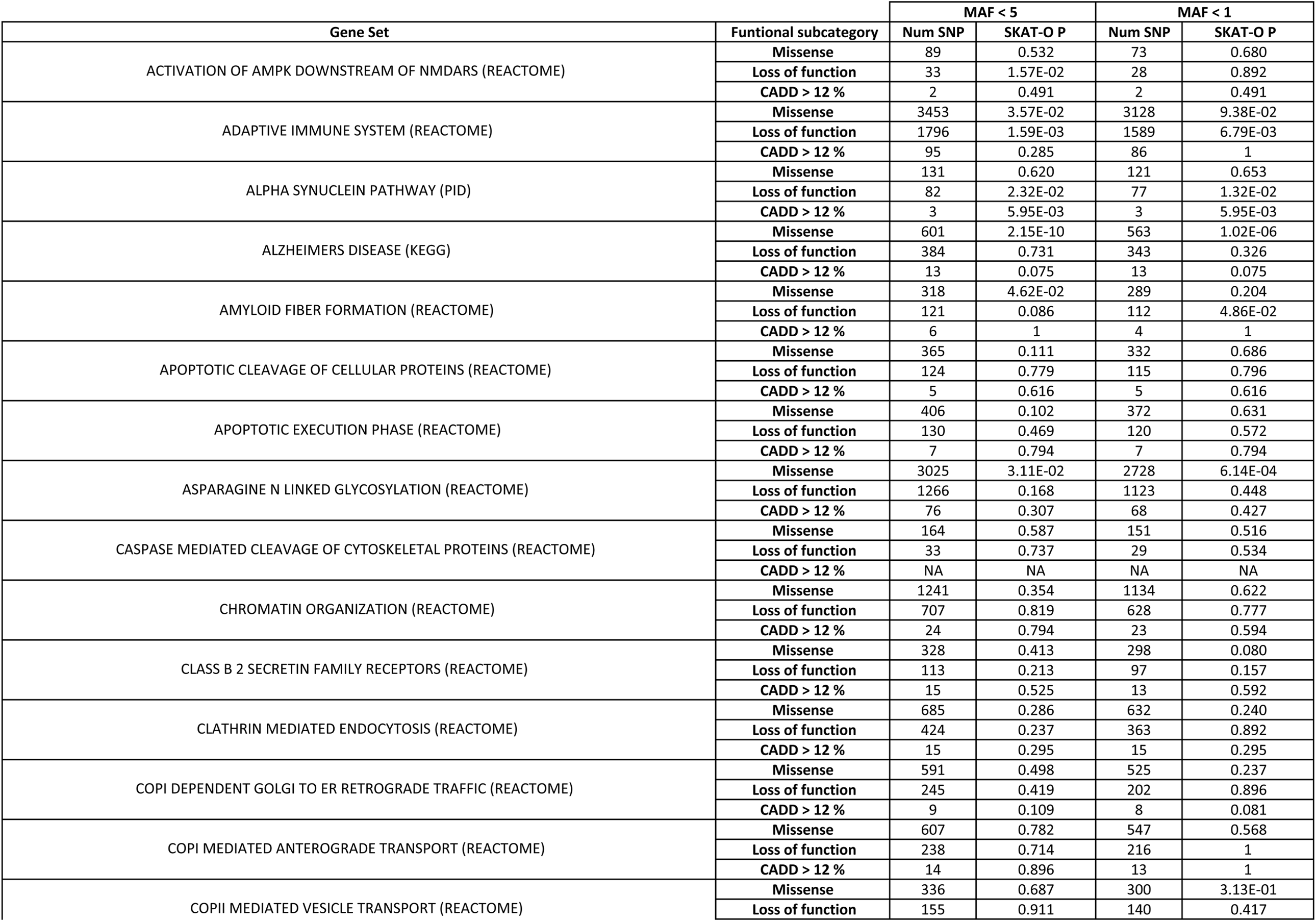
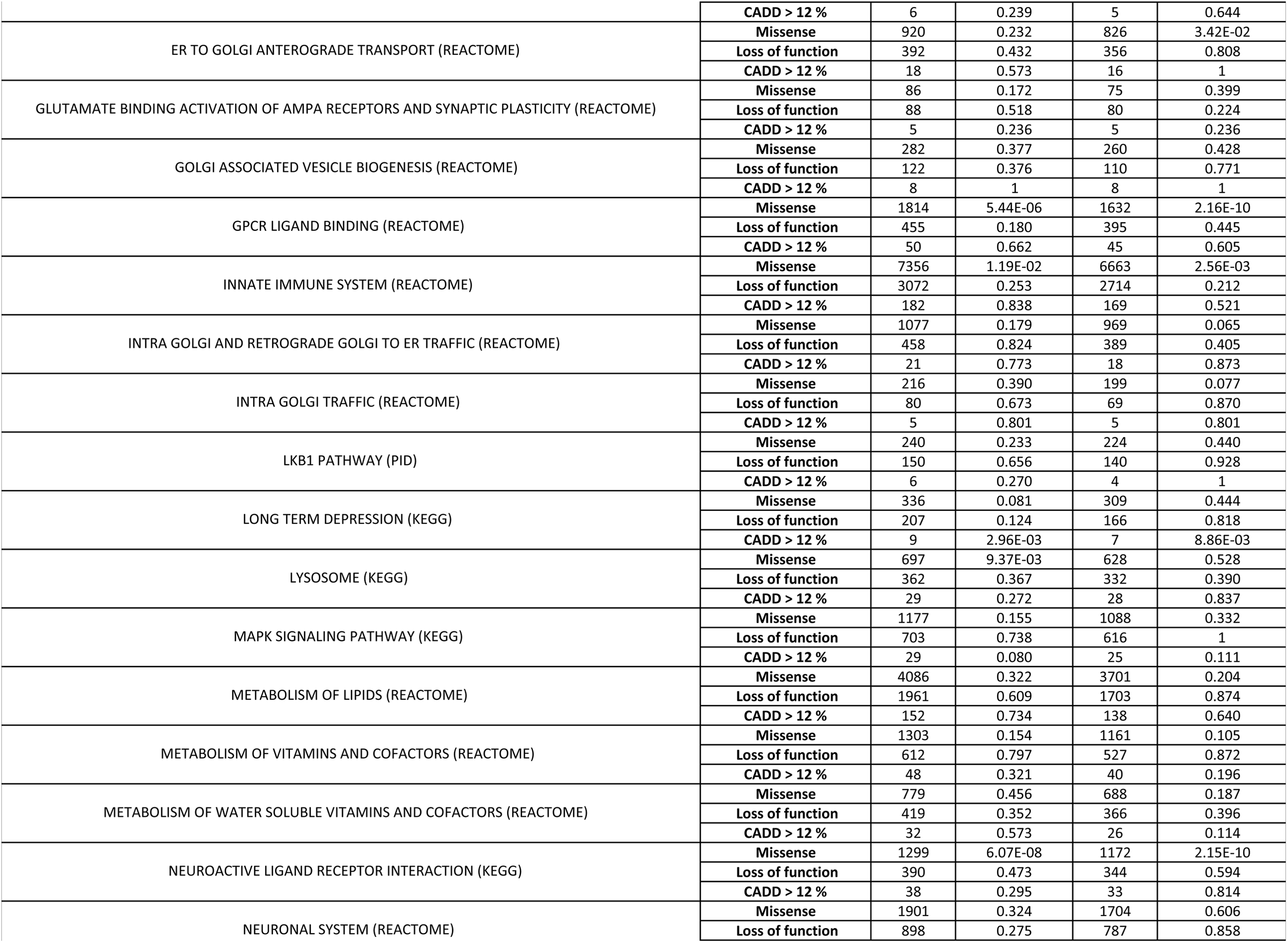
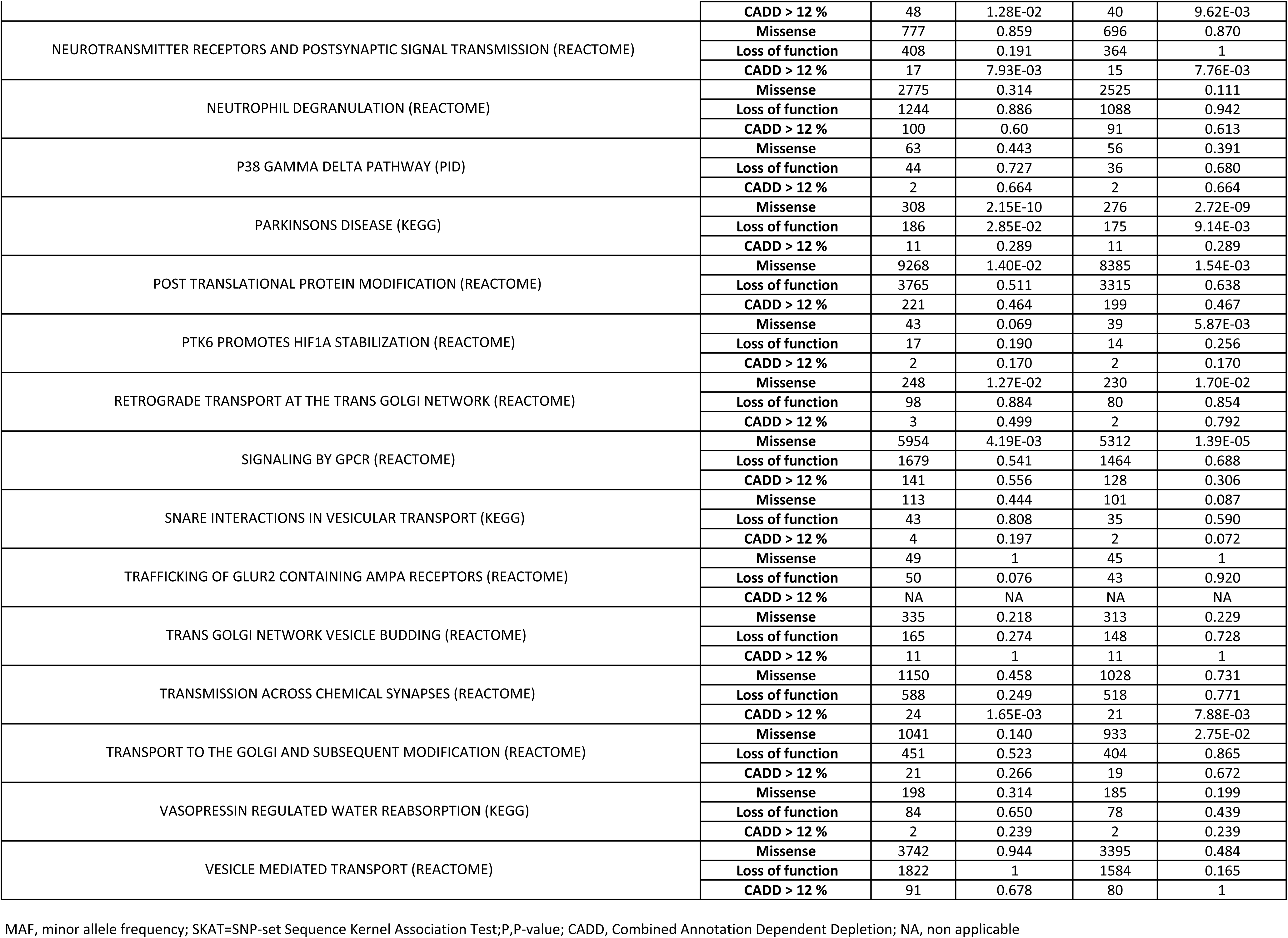
Association of canonical pathways and PD risk through rare variation.

After removing PD GWAS hits and SNPs located 1Mb upstream and downstream, innate immune system and signaling by GPCR remained significantly associated with PD suggesting that rare variation within these pathways contributes to PD heritability **(Supplementary Table 4)**.

### Transcriptome map reveals expression modules linked to PD etiology

Using Louvain community detection, we generated transcriptomic networks among PD cases. We identified 54 d*e novo* expression communities **(Supplementary Table 5, Supplementary Figure 3).** Overall, the communities generated were relatively robust, with a modularity score of 0.523 (modularity ranges from -1 to 1, with closer to 1 suggesting stronger connectivity between network members). The 54 network communities were found to be enriched via hypergeometric tests after Bonferroni correction for processes relating to immune system response, ribosome RNA processing to the nucleus and cytosol, cell cycle, oxidative stress, and mitochondrial impairment **(Figure 3, Supplementary Table 6).**

**Figure 3.**
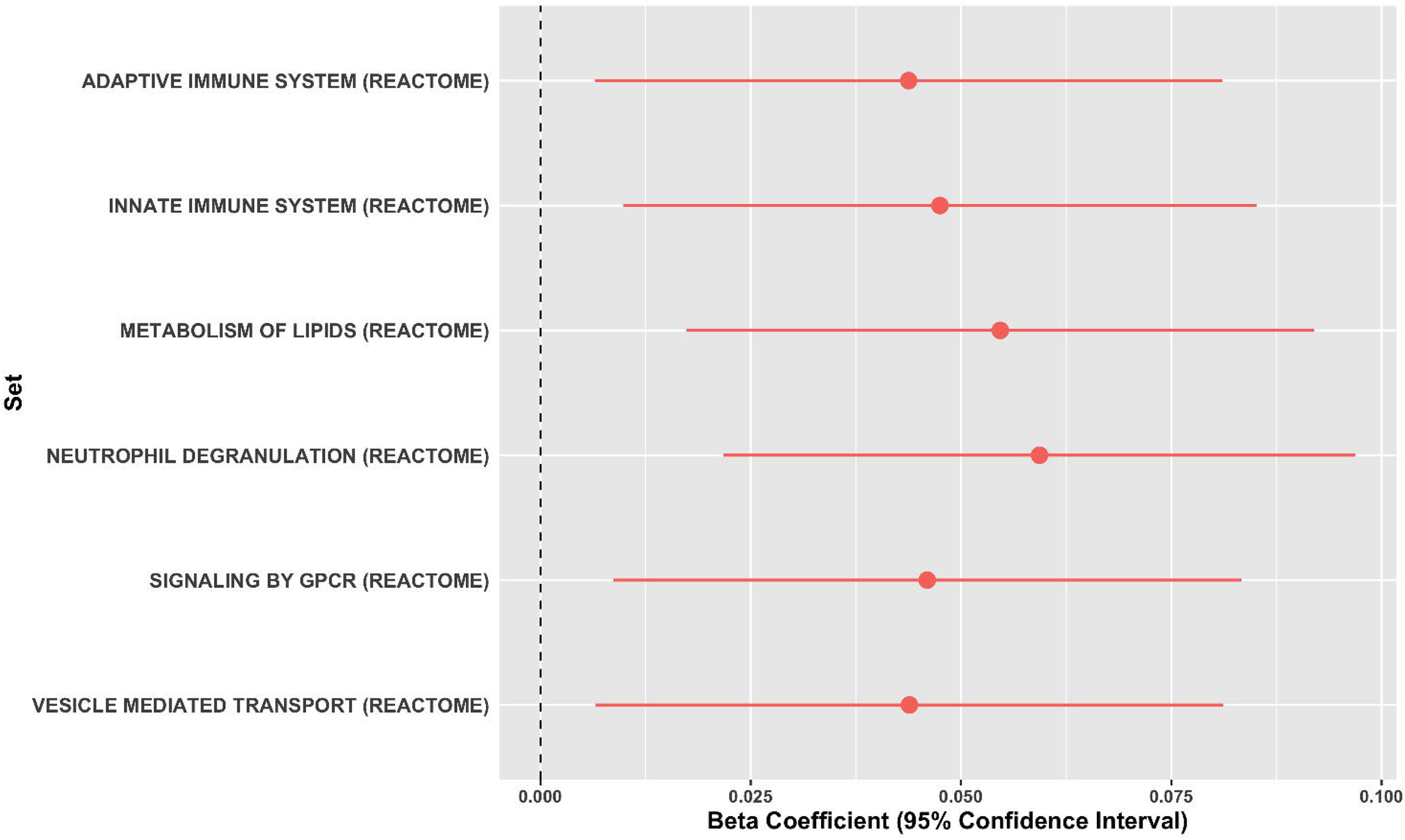
Functional enrichment analyses of transcriptomic community maps. The x-axis represents the gene set enrichment (%) based on the community map gene lists. Intersection size denotes the number of input genes within an enrichment category. Blue color indicates the adjusted association p-values on a -log10 scale. *** (by chemiosmotic coupling and heat production by uncoupling proteins).

### Dopaminergic adult neurons, serotonergic neurons and radial glia play a role on PD etiology

We used single-cell RNA sequencing data from 24 different brain cell types (Skene et al., 2018) [31]. For each of those cell types, genes were clustered into 5 gene sets according to the level of expression specificity, ranging from 0 to 1 (0 means that a gene is not expressed at all and 1 means the gene expression is highly specific for that cell type). Then, PRS was calculated per quintile of specificity within cells. Increased PRS R^2^, consistent with increased cell expression specificity, was observed for dopaminergic adult neurons, serotonergic neurons and radial glia at P < 0.05 in both the *training* and *replication phases* (**Supplementary Table 7, Supplementary Figure 4**, https://pdgenetics.shinyapps.io/pathwaysbrowser/).

### Mendelian Randomization prioritizes pathways and genes based on their functional consequence

We aimed at nominating genes within significant pathways contributing to PD etiology by assessing changes in expression across blood and brain. Out of the 46 pathways of interest, seven showed a significant enrichment of QTLs more than expected by chance in the brain, one in substantia nigra and eleven in blood **(Supplementary Table 8)**. SMR revealed functional genomic associations with expression QTLs in 201 genes (**Supplementary Table 9**) of which 88 were found to be part of the network communities significantly associated with PD in our transcriptome community map **(Supplementary Table 10)**.

## DISCUSSION

Despite success at uncovering genetic risk factors associated with PD, our understanding of the molecular processes involved in disease is still limited. Using the largest genomic and transcriptomic PD cohorts currently available, our study sought to define both cumulative genetic risk and functional consequences linked to myriad biological pathways in an unbiased and data-directed manner. To our knowledge, there are no previous reports in the PD field where a similar approach has been implemented to explore the contribution of thousands of molecular processes on both the trigger (risk) and the effect (expression changes) in a systematic manner.

Our large-scale PRS analysis identified multiple biological pathways associated with PD risk through common genetic variation. Overall, our results found that molecular processes underlying protein misfolding and aggregation, post-translational protein modification, immune response, membrane and intracellular trafficking, lipid metabolism, synaptic transmission, endosomal-lysosomal dysfunction and apoptosis mediated by initiator and executioner caspases are among the main contributors to PD etiology.

PD heritability remains incompletely deciphered by the genes and variants identified to date [15]. Here we demonstrate that some of these significant pathways contribute to the heritability of PD outside of what is explained by current GWAS [15]. Notably, our genetic analyses provide definitive evidence for the role of several signal transduction mechanisms affecting adaptive and innate immune response, vesicular mediated transport, and lipid metabolism on the risk for PD even after excluding PD known GWAS loci. The present study suggests that additional targets within these pathways are yet to be identified and prioritizes genes for follow-up functional studies.

A novel aspect of our study is that we nominate pathways whose implication on PD pathology has been poorly studied or debatable before. Our results support the hypothesis that chromatin remodeling and epigenetic mechanisms contribute to the development of PD [11]. An appropriate balance and distribution of active and repressed chromatin is required for proper transcriptional control, maintaining nuclear architecture and genomic stability, as well as regulation of the cell cycle [8]. Dysfunction in the epigenetic machinery has been shown to play a role in the etiology of a number of neurodegenerative and neurodevelopmental disorders either by genetic variation in an epigenetic gene or by changes in DNA methylation or histone modifications [11]. Similarly, our approach supports a role for vitamin metabolism on PD risk. Vitamins are crucial cofactors in the metabolism of carbohydrates, fat and proteins, and vitamin deficiency has been widely proven to promote oxidative stress and neuroinflammation [14].

Interestingly, some of the nominated pathways seem to span the etiological risk spectrum in which both common and rare variation contribute to PD susceptibility. In concordance with previous studies [19], our study identified an increased collective effect of rare lysosomal related variants in PD etiology. Additionally, we found evidence for a burden of rare damaging alleles in a range of specific processes, including neuronal transmission-related pathways and immune response.

The present study represents a significant step forward in our understanding of important connections between genetic factors, functional consequences and PD etiology. We constructed a transcriptome map by clustering *de novo* pathways relevant to disease pathology. Functional characterization analysis of these expression communities revealed that dysregulation of the immune system and inflammatory response including neutrophil degranulation, interferon alpha beta signaling and other cytokine-related signaling pathways are key disease processes. Strikingly, when looking at molecular mechanisms significantly associated with PD risk, a cumulative effect of rare loss of function variants was found to be linked to disease through the adaptive immune system pathway. Both inflammation and autoimmune response have been widely studied with regard to PD etiology. Previous genetic studies have identified risk loci spanning key immune-associated genes such as *BST1 (bone marrow stromal cell antigen 1)*, a gene known to play role in neutrophil adhesion and migration, and *HLA* (human leukocyte antigen) [21] [15]. In support of this, it has been reported that α-synuclein-derived fragments act as antigenic epitopes displayed by HLA receptors, where both helper and cytotoxic T-cell responses are present in a high percentage of patients when tested [25].

Our analysis provides compelling evidence that dysregulation in genes that play a pivotal role in mitochondrial homeostasis exists in genetically complex PD. Despite not identifying these pathways as part of the stringent large-scale PRS analysis, our transcriptome community map showed an enrichment for the respiratory electron transport ATP synthesis by chemiosmotic coupling process and mitochondrial oxidative phosphorylation, in concordance with other reports [33]. Among the expression networks to highlight, it should be pointed out an enrichment in cell cycle and cell death machinery related processes and ribosome RNA processing to the nucleus and cytosol.

Our study aimed at pinpointing the specific drivers underlying these significant networks. Focusing on pathways linked to PD risk, SMR was applied to prioritize genes whose variation was found to be associated with expression changes linked with PD risk. Interestingly, we managed to replicate 88 of these genes after validating the functional consequence within our transcriptome community map.

Despite genetic efforts, it remains a matter of study in what cell types risk variants are active, which is essential for understanding etiology and experimental modeling. By integrating genetics and single-cell expression data, we found that PD risk is linked to expression specificity patterns in dopaminergic neurons, serotonergic neurons, and radial glia, suggesting that these cell types disrupt biological networks that impact PD risk. Although our study failed at replicating specific enrichment patterns for oligodendrocytes and microglia as previously reported using other approaches [18], our results are in concordance with previous literature that applies various methodologies to gain similar conclusions [5].

The strengths of this study include an unbiased effort to link risk variants to biological pathways and characterize the functional consequence. While this study marks major progress in integrating human genetic and functional evidence, much remains to be established. A caveat of this study is that our approach was limited by the canonical gene sets publicly defined that were used for pathway analysis, and the relatively few brain regions studied for cell type analysis, which was based on mice data. We are aware that additional molecular networks and cell types from unsampled regions could contribute to PD. In addition, PRS analyses considered that all the variants conferred risk under the additive model and did not cover regulatory regions adjacent to the up or downstream of the genes or intergenic variants, that may be crucial for disease. A further limitation of our study is that although we used state-of-the-art methodologies such as SMR to nominate candidate pathways and genes related to PD etiology, QTL datasets and associations are affected by both small sample size and low cis-SNP density. In addition, trans-QTL could not be assessed. Furthermore, our study focused on individuals of European ancestry, given that large sample sizes were required to create this resource. Replication in ancestrally diverse populations would be necessary for future studies. We anticipate that substantial collaborative efforts will lead to an improvement in statistical power and accuracy to define pathways linked to PD.

In conclusion, our high-throughput and hypothesis-free approach exemplifies a powerful strategy to provide valuable mechanistic insights into PD etiology and pathogenesis. We highlight several promising pathways, cell types, and genes for further functional prioritization, aware that further in-depth investigation will be required to prove a definite link. As part of this study, we created a foundational resource for the PD community that can be applied to other neurodegenerative diseases with complex genetic etiologies (https://pdgenetics.shinyapps.io/pathwaysbrowser/). In future studies, linking specific phenotypic aspects of PD to pathways will constitute a critical effort by using large longitudinal cohorts of well clinically characterized PD patients, with the hope of yielding disease-modifying therapeutic targets that are effective across PD subtypes.

## Funding agencies

This research was supported in part by the Intramural Research Program of the National Institutes of Health (National Institute on Aging, National Institute of Neurological Disorders and Stroke; project numbers: project numbers 1ZIA-NS003154, Z01-AG000949-02 and Z01-ES101986). In addition, this work was supported by the Department of Defense (award W81XWH-09-2-0128), and The Michael J Fox Foundation for Parkinson’s Research. C.R.S. was supported in part by NIH grants U01NS095736, U01NS100603, R01AG057331, and R01NS115144 and by the MJFF.

## ACKNOWLEDGMENTS

We would like to thank all of the subjects who donated their time and biological samples to be a part of this study. This work was supported in part by the Intramural Research Programs of the National Institute of Neurological Disorders and Stroke (NINDS), the National Institute on Aging (NIA), and the National Institute of Environmental Health Sciences both part of the National Institutes of Health, Department of Health and Human Services; project numbers Z01-AG000949-02 and Z01-ES101986. In addition this work was supported by the Department of Defense (award W81XWH-09-2-0128), and The Michael J Fox Foundation for Parkinson’s Research. We would also like to thank all members of the International Parkinson Disease Genomics Consortium (IPDGC). For a complete overview of members, acknowledgements and funding, please see http://pdgenetics.org/partners. We would like to thank the Accelerating Medicines Partnership initiative (AMP-PD) for the publicly available whole genome sequencing data.

This work utilized the computational resources of the NIH HPC Biowulf cluster. (http://hpc.nih.gov) We thank contributors who collected samples used in this initiative, as well as patients and families, whose help and participation made this work possible. We thank members of the North American Brain Expression Consortium (NABEC) for providing DNA samples derived from brain tissue. Brain tissue for the NABEC cohort was obtained from the Baltimore Longitudinal Study on Aging at the Johns Hopkins School of Medicine, and from the NICHD Brain and Tissue Bank for Developmental Disorders at the University of Maryland, Baltimore, MD, USA. We would like to thank the United Kingdom Brain Expression Consortium (UKBEC) for providing DNA samples. This study acknowledges the National Institute of Neurological Disorders and Stroke (NINDS) supported Parkinson’s Disease Biomarkers Program Investigators (https://pdbp.ninds.nih.gov/sites/default/files/assets/PDBP_investigator_list.pdf). A full list of PDBP investigators can be found at https://pdbp.ninds.nih.gov/policy. Data and biospecimens used in the preparation of this manuscript were obtained from the Parkinson’s Disease Biomarkers Program (PDBP) Consortium, part of the National Institute of Neurological Disorders and Stroke at the National Institutes of Health. Investigators include: Roger Albin, Roy Alcalay, Alberto Ascherio, Thomas Beach, Sarah Berman, Bradley Boeve, F. DuBois Bowman, Shu Chen, Alice Chen-Plotkin, William Dauer, Ted Dawson, Paula Desplats, Richard Dewey, Ray Dorsey, Jori Fleisher, Kirk Frey, Douglas Galasko, James Galvin, Dwight German, Lawrence Honig, Xuemei Huang, David Irwin, Kejal Kantarci, Anumantha Kanthasamy, Daniel Kaufer, James Leverenz, Carol Lippa, Irene Litvan, Oscar Lopez, Jian Ma, Lara Mangravite, Karen Marder, Laurie Ozelius, Vladislav Petyuk, Judith Potashkin, Liana Rosenthal, Rachel Saunders-Pullman, Clemens Scherzer, Michael Schwarzschild, Tanya Simuni, Andrew Singleton, David Standaert, Debby Tsuang, David Vaillancourt, David Walt, Andrew West, Cyrus Zabetian, Jing Zhang, and Wenquan Zou. The PDBP Investigators have not participated in reviewing the data analysis or content of the manuscript. This work was supported by Scripps Research Translational Institute, an NIH-NCATS Clinical and Translational Science Award (CTSA; 5 UL1 RR025774). We are grateful to the NIH NeuroBioBank for providing brain tissue samples for Parkinson’s disease cases. We are grateful to the Banner Sun Health Research Institute Brain and Body Donation Program of Sun City, Arizona, for the provision of human brain tissue (PI: Thomas G. Beach, MD). The Brain and Body Donation Program is supported by the National Institute of Neurological Disorders and Stroke (U24 NS072026 National Brain and Tissue Resource for Parkinson’s Disease and Related Disorders), the National Institute on Aging (P30 AG19610 Arizona Alzheimer’s Disease Core Center), the Arizona Department of Health Services (contract 211002, Arizona Alzheimer’s Research Center), the Arizona Biomedical Research Commission (contracts 4001, 0011, 05-901 and 1001 to the Arizona Parkinson’s Disease Consortium) and the Michael J. Fox Foundation for Parkinson’s Research.

## AUTHORS’ ROLES

1. **Research project:** A. Conception (SBC, SSA, MN, CD, AS), B. Organization (SBC, SSA, MN), C. Execution (SBC, SSA, JK, MM, MN)
2. **Data generation:** A. Experimental (RG, JD, CB, MN).
3. **Statistical Analysis:** A. Design (SBC, SSA,MN), B. Execution (SBC, SSA, MM, MN)
4. **Manuscript Preparation:** A. Writing of the first draft (SBC), B. Review and Critique (all authors)

## FIGURES

**Supplementary Figure 1.**
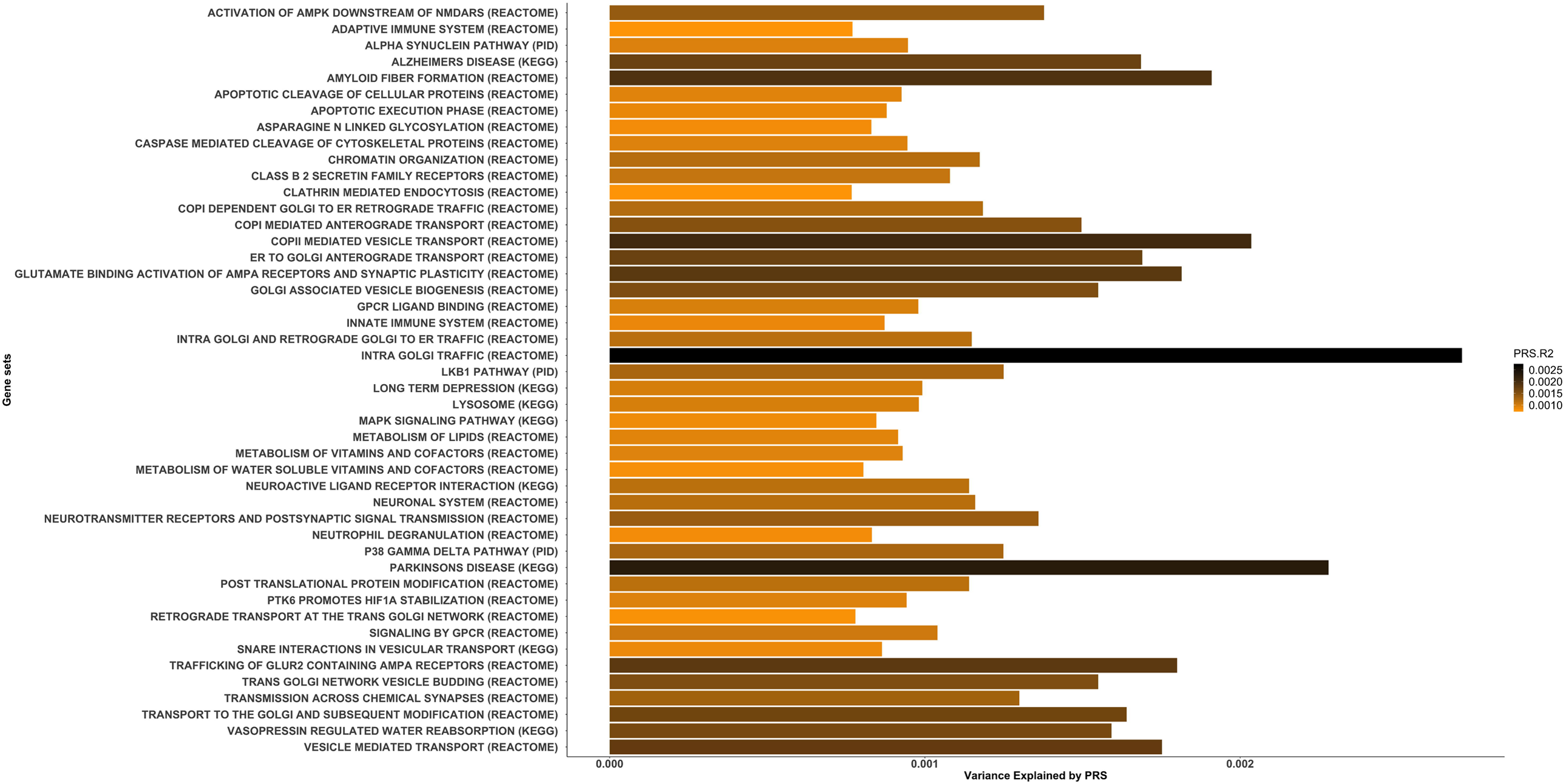
Canonical Pathways associated with PD risk clustered in hierarchies according to genetic redundancy

**Supplementary Figure 2.**
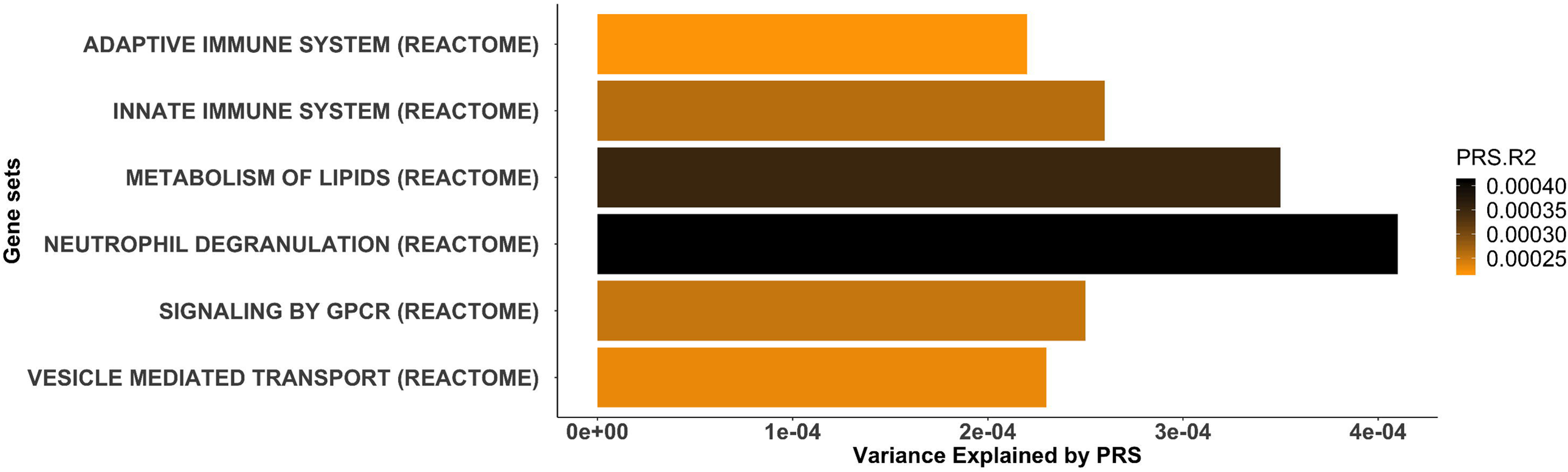
Canonical Pathways associated with PD risk clustered in hierarchies according to published literature and public curated databases

**Supplementary Figure 3.**
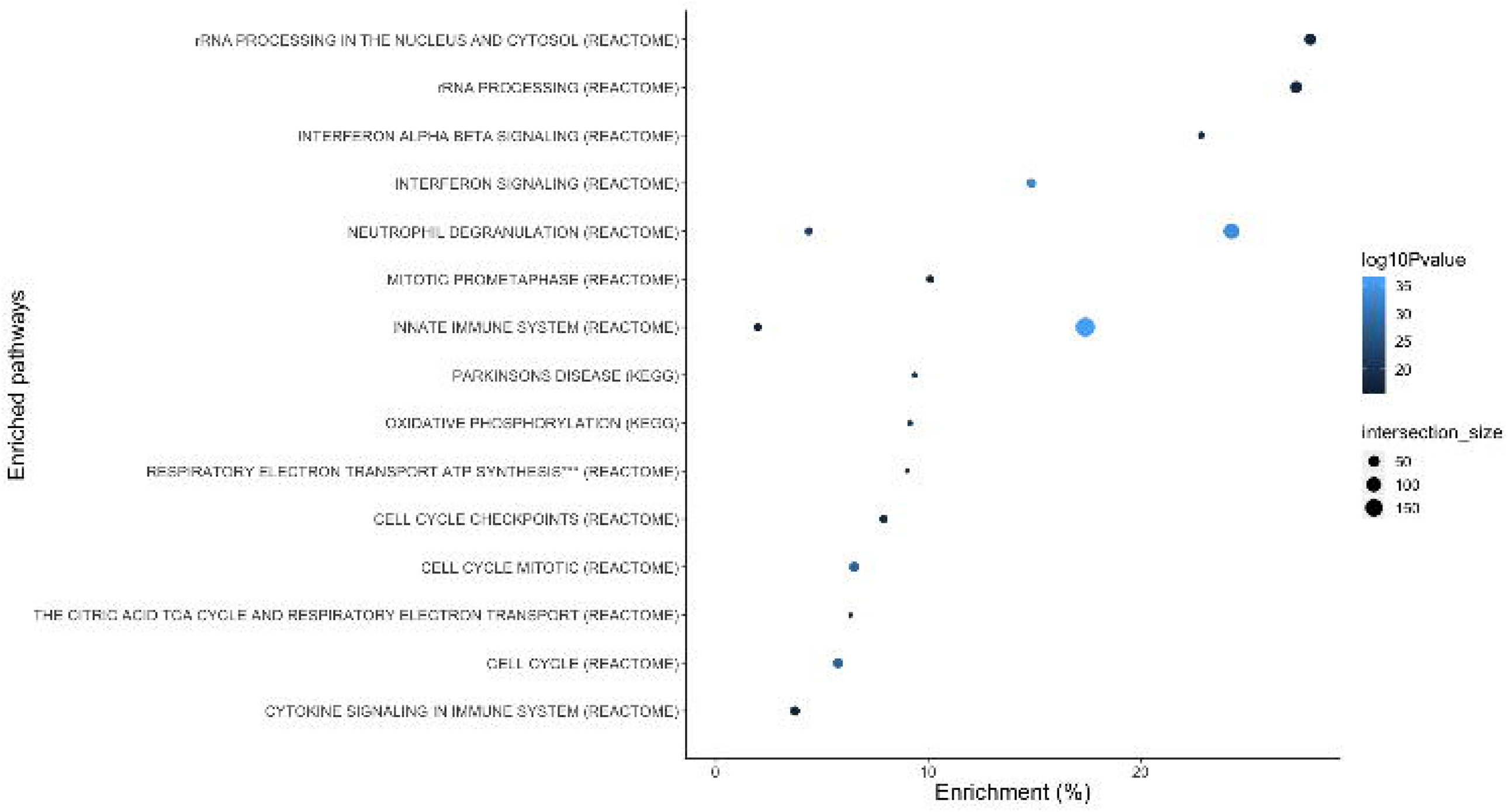
Distribution of transcriptomic community networks according to Louvain cluster detection. Colors represent different transcriptomic communities. Circles represent genes within expression networks.

**Supplementary Figure 4. Cell types significantly associated with PD etiology based on expression specificity patterns.** AE, astrocytes ependymal; Dopamin, dopaminergic; E., embryonic; Nu, nucleus; Lept, Leptomeningeal; MSN, medium spiny neurons; Oxyt.&Vasop., Oxytocin and Vasopressin Expressing Neurons;P, precursor;Hypoth, hypothalamic;GABA, GABAergic;GLUT, Glutamatergic.

## TABLES

**Supplementary Table 1. Demographic and clinical characteristics of the GWAS cohorts under study**

**Supplementary Table 2. Demographic and clinical characteristics of the WGS cohorts under study.**

**Supplementary Table 3. Canonical pathways significantly associated with PD risk in the discovery phase**

**Supplementary Table 4. Gene-set burden analyses of canonical pathways significantly associated with PD risk excluding PD known risk loci +/- 1Mb upstream and downstream**

**Supplementary Table 5. Genes within expression community maps and modularity ranges**

**Supplementary Table 6. Functional enrichment analysis of network community maps**

**Supplementary Table 7. Cell types significantly associated with PD etiology based on expression specificity enrichment analyses**

**Supplementary Table 8. Functional enrichment for expression quantitative trait loci across pathways significantly associated with PD risk**

**Supplementary Table 9. Functional associations for genes within significant pathways associated with PD risk by two sample Mendelian randomization**

**Supplementary Table 10. Genes showing functional inferences through two sample Mendelian randomization and functional consequence in the transcriptome communities International Parkinson’s Disease Genomics Consortium members (IPDGC)**

