## Supplementary material for "Large-scale pathway-specific polygenic risk, transcriptomic community networks and functional inferences in Parkinson disease": Suplementary information: Sup_Figures.pdf

**Supplementary Figure 1. Canonical Pathways associated with PD risk clustered in hierarchies according to genetic redundancy**

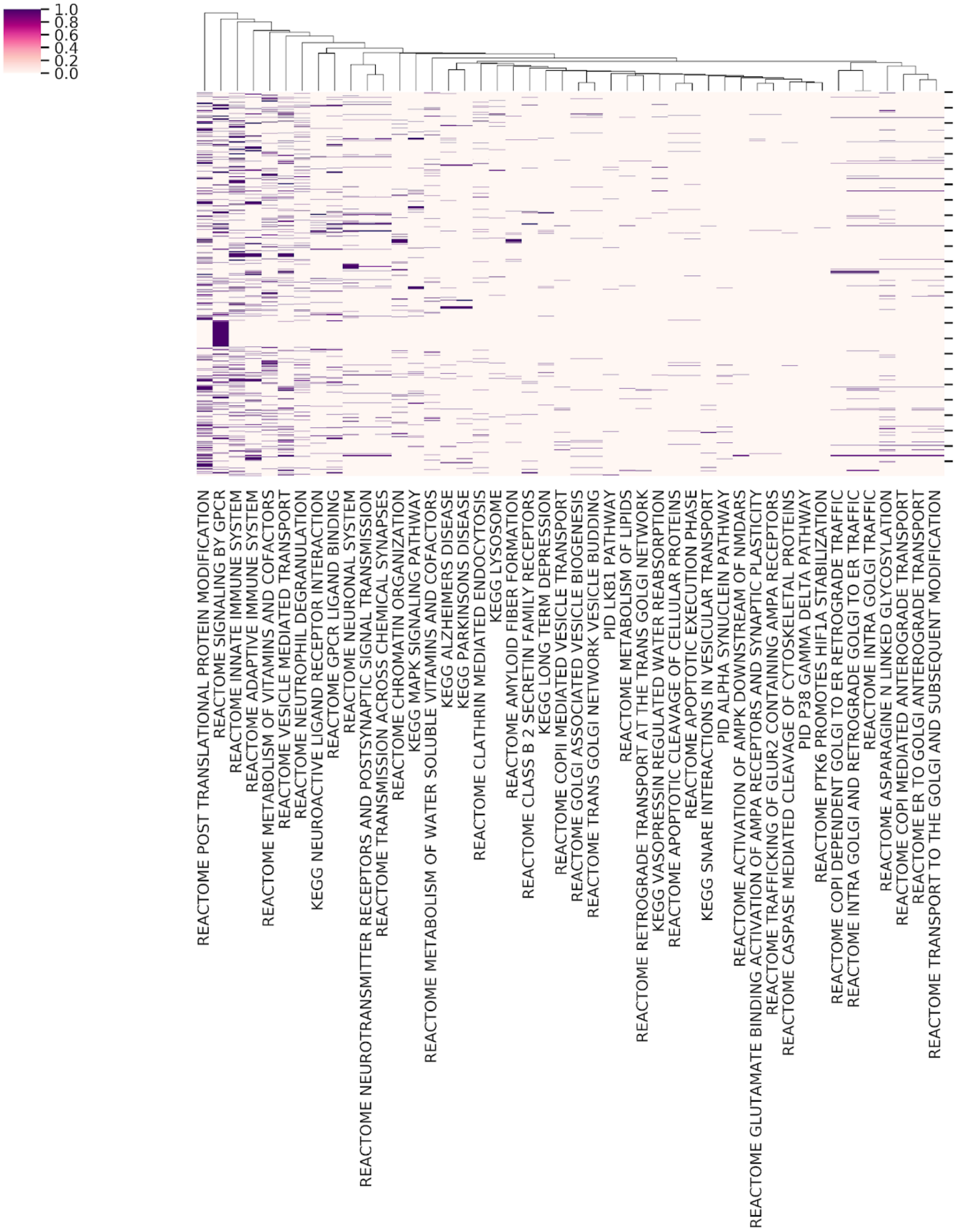

**Supplementary Figure 2. Canonical Pathways associated with PD risk clustered in hierarchies according to published literature and public curated databases**

**Metabolism**

**Protein metabolism**

- Alpha synuclein pathway
- Amyloid fiber formation
- Asparagine N-linked glycosylation
- Post-translational protein modification
- Transport to the Golgi and subsequent modification

**Lipid metabolism**

**Metabolism of vitamins and cofactors**

**Programmed cell death**

- Apoptotic cleavage of cellular proteins
- Apoptotic execution phase
- Caspase mediated cleavage of cytoskeletal proteins

**Immune system**

- Adaptive immune system
- Innate immune system
- Neutrophil degranulation

**Vesicle-mediated transport and membrane trafficking**

- Clathrin-mediated endocytosis
- COPI-dependent Golgi-to-ER retrograde traffic
- COPI-mediated anterograde transport
- COPII-mediated vesicle transport
- ER-to-Golgi anterograde transport
- Golgi-associated vesicle biogenesis
- Intra-Golgi and retrograde Golgi-to-ER
- Lysosome
- Retrograde transport at the trans-Golgi network
- SNARE interactions in vesicular transport
- Trans-Golgi network vesicle budding

**Signal transduction**

- Class B-2 secreting family receptors
- GPCR signaling and ligand binding
- LKB1 pathway
- MAPK signaling pathway
- Neuroactive ligand receptor interaction
- P38 gamma delta pathway
- PTK6 promotes HIF1A stabilization

**Neuronal transmission across chemical synapses**

- Activation of AMPK downstream of NMDA receptors
- Glutamate binding activation of AMPA receptors and synaptic plasticity
- Neurotransmitter receptors and postsynaptic signal transmission
- Trafficking of GLUR2 containing AMPA receptors
- Transmission across chemical synapses

**Miscellaneous**

- Chromatin organization
- Alzheimer disease
- Parkinson disease
- Vasopressin-regulated water reabsorption
- Long-term depression

**Supplementary Figure 3. Distribution of transcriptomic community networks according to Louvain cluster detection**

Colors represent different transcriptomic communities. Circles represent genes within expression networks.

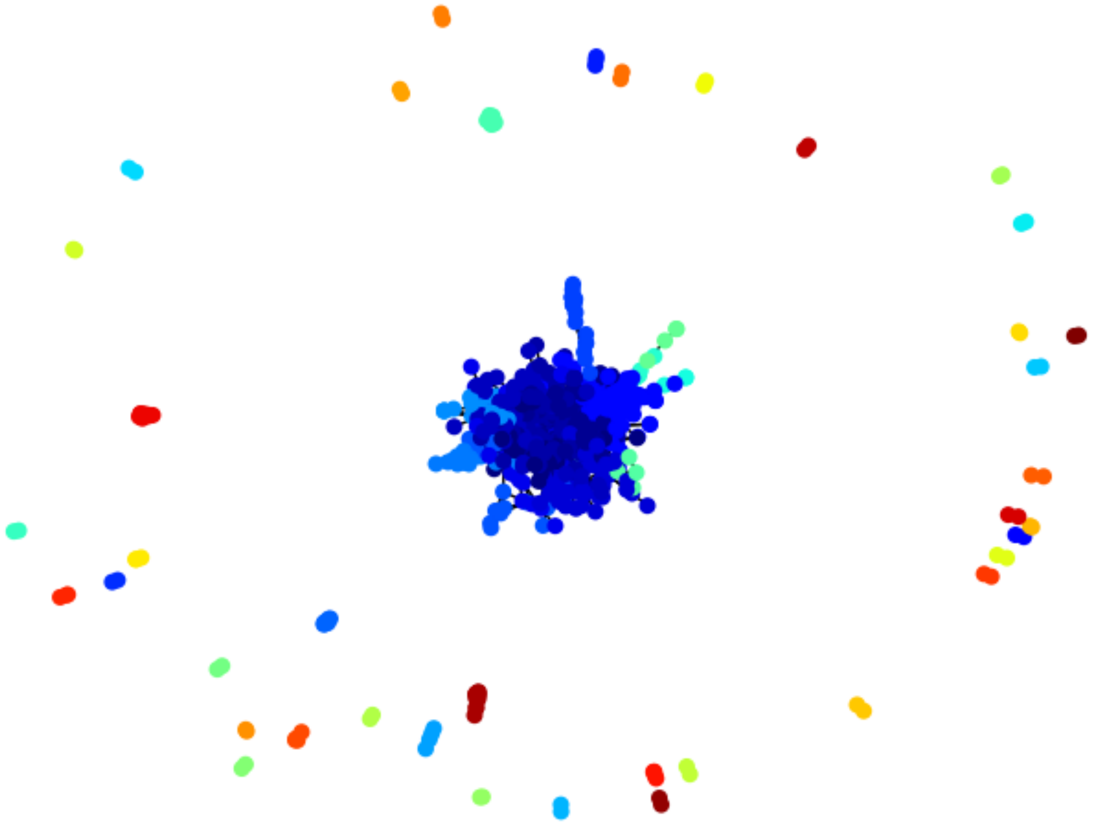

**Supplementary Figure 4. Cell types significantly associated with PD etiology based on expression specificity patterns.**

AE, astrocytes ependymal; Dopamin, dopaminergic; E., embryonic; Nu, nucleus; Lept, Leptomeningeal; MSN, medium spiny neurons; Oxyt.&Vasop., Oxytocin and Vasopressin Expressing Neurons;P, precursor;Hypoth, hypothalamic;GABA, GABAergic;GLUT, Glutamatergic.

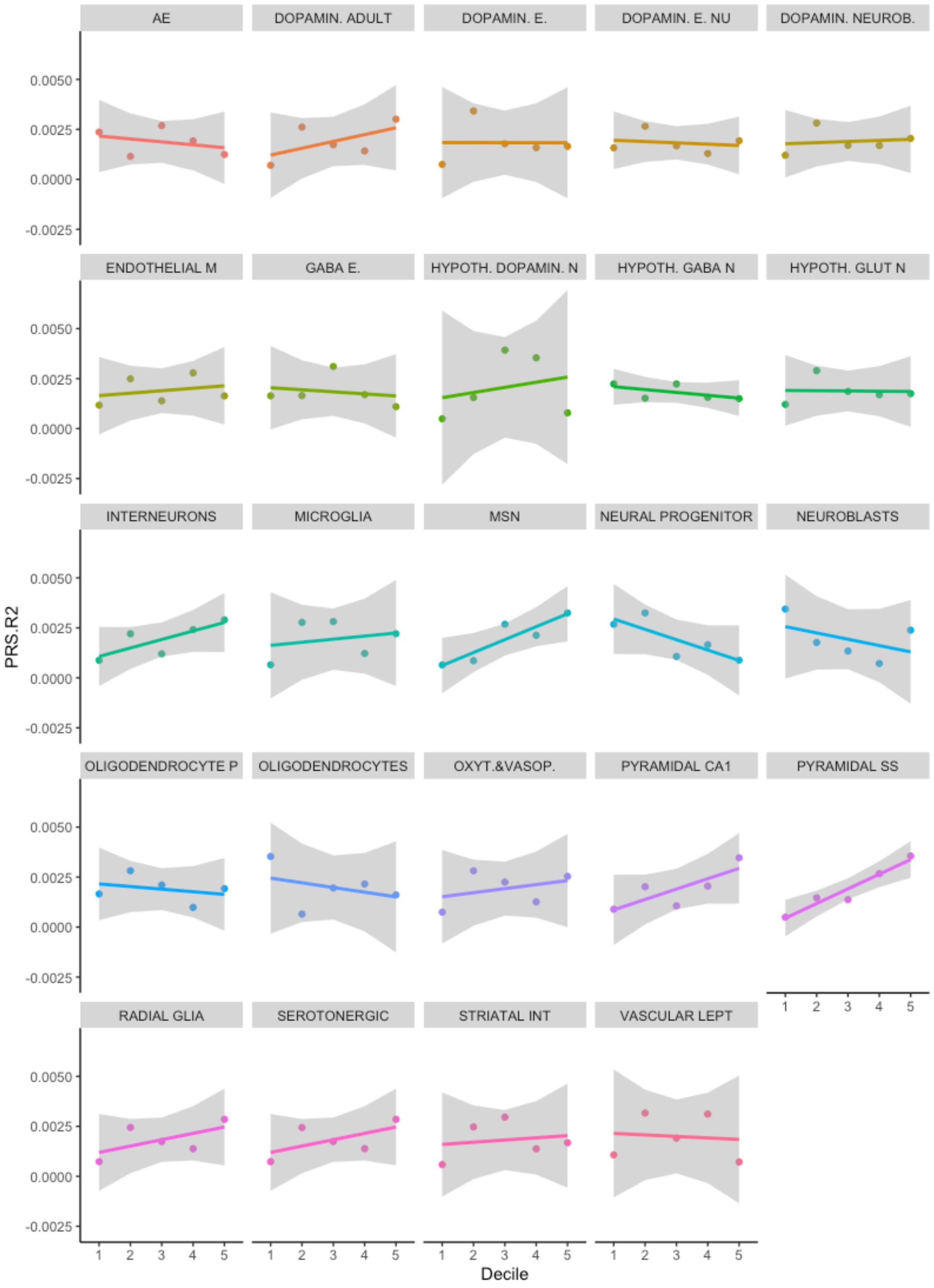
